# Using AnnoTree to get more assignments, faster, in DIAMOND+MEGAN microbiome analysis

**DOI:** 10.1101/2021.11.23.469797

**Authors:** Anupam Gautam, Hendrik Felderhoff, Caner Bağci, Daniel H. Huson

## Abstract

In microbiome analysis, one main approach is to align metagenomic sequencing reads against a protein-reference database such as NCBI-nr, and then to perform taxonomic and functional binning based on the alignments. This approach is embodied, for example, in the standard DIAMOND+MEGAN analysis pipeline, which first aligns reads against NCBI-nr using DIAMOND and then performs taxonomic and functional binning using MEGAN. Here we propose the use of the AnnoTree protein database, rather than NCBI-nr, in such alignment-based analyses to determine the prokaryotic content of metagenomic samples. We demonstrate a 2-fold speedup over the usage of the prokaryotic part of NCBI-nr, and increased assignment rates, in particular, assigning twice as many reads to KEGG. In addition to binning to the NCBI taxonomy, MEGAN now also bins to the GTDB taxonomy.

**IMPORTANCE:** The NCBI-nr database is not explicitly designed for the purpose of microbiome analysis and its increasing size makes its unwieldy and computationally expensive for this purpose. The AnnoTree protein database is only one quarter the size of the full NCBI-nr database and is explicitly designed for metagenomic analysis, and so should be supported by alignment-based pipelines.

## INTRODUCTION

Next-generation sequencing (NGS) has revolutionized many areas of biological research (1, 2), providing ever-more data at an ever-decreasing cost. One such area is microbiome research, the study of microbes in their theater of activity using metagenomic sequencing (3). Here, deep short-read sequencing, and improving performance of long-read sequencing, are helping to explore the roles and interactions of microbiomes in different environments.

The two initial tasks for any microbiome sequencing dataset are (1) taxonomic analysis, that is, to determine which organisms are present in a microbiome sample, and (2) functional analysis, to determine which genes and pathways are present. Task (1) can be addressed, to a degree, using amplicon sequencing (targeting 16S rRNA, 18S rRNA or ITS, say). However, a species- or strain-level taxonomic analysis, and task (2), both are best addressed using whole-genome shotgun sequencing, that is, “metagenomics proper”.

Metagenomic shotgun sequencing reads can be analyzed using a number of different approaches, such as alignment-free *k*-mer analysis (4, 5, 6), alignment against DNA references (7, 8) or translated-alignment against protein references (9, 10, 11, 12, 13).

The DIAMOND+MEGAN approach uses DIAMOND (14) to perform translated alignment (short-read sequencing), or “frameshift-aware” translated alignment (long-read sequencing) of metagenomic reads or contigs against the NCBI-nr database (15). The resulting alignment files are then “meganized” or analyzed using MEGAN (16) so as to perform taxonomic and functional binning. A detailed protocol of this simple two-step pipeline is presented in (17).

During meganization, reads are assigned to taxonomic classes in the NCBI taxonomy (18) and, now also to the GTDB taxonomy (19). Reads are also assigned to functional entities in EC (20), EggNOG (21), InterPro families (22), KEGG (23) and SEED (24, 25).

The DIAMOND+MEGAN pipeline was originally designed for use with the NCBI-nr database (10). The NCBI-nr database contains non-identical protein sequences from GenBank CDS translations, PDB (26), Swiss-Prot (27), PIR, and PRF, covering all domains of life, and viruses. In August 2021, NCBI-nr comprised over 420 million sequences, of which just over half, ≈ 213 million, belong to bacteria or archaea. The NCBI-nr database is not explicitly designed for use in metagenomic-analysis and its rapidly-increasing size is making it unwieldy for this purpose.

To perform taxonomic binning, MEGAN assigns reads to the NCBI taxonomy (28, 18), which contains more than 2.2 million nodes and is designed for “structuring communication concerning all forms of life on Earth”. More recently, the GTDB taxonomy (19) was developed for the explicit purpose of taxonomic analysis of microbiome sequencing data. Here, we introduce support for the GTDB taxonomy in MEGAN. Now, the DIAMOND+MEGAN pipeline performs taxonomic binning of metagenomic reads according to both the NCBI taxonomy and the GTDB taxonomy.

AnnoTree (29) provides functional annotations from over 27,000 bacterial and 1,500 archaeal genomes obtained from the GTDB database (19). The total number of protein sequences contained in the AnnoTree database is 106, 052, 079. The AnnoTree database is approximately one quarter the size of NCBI-nr and only half the size of the prokaryotic part of NCBI-nr. However, AnnoTree contains protein sequences from more metagenome assembled genomes (MAGs) than NCBI-nr, and these are more-likely to be found in microbiome samples.

In this paper, we describe and provide the necessary files for performing DIAMOND+MEGAN analysis using the AnnoTree protein reference database, as an alternative to NCBI-nr, to analyze the prokaryotic content of metagenomic sequencing samples. To illustrate the utility of this approach, first we compare the performance of DIAMOND+MEGAN using NCBI-nr and AnnoTree protein-reference sequences on a published mock-community of 23 bacterial and 3 archaeal strains (30). Then, using ten published datasets from a range of different environments and obtained using both short- and long-read sequencing techniques, we demonstrate that AnnoTree-based analysis is twice as fast and has a higher assignment-rate than NCBI-nr-based analysis, when using the DIAMOND+MEGAN pipeline. We also compare examples of both the NCBI and GTDB taxonomic binning.

## RESULTS

Using the standard DIAMOND+MEGAN analysis pipeline (16), metagenomic sequencing reads are first aligned against the NCBI-nr database using DIAMOND and then the resulting alignments are processed by MEGAN (or the command-line tool daameganizer) so as to perform taxonomic and functional binning. We will refer to this as an “NCBI-nr run” of the pipeline.

In this paper we describe a new application of the DIAMOND+MEGAN pipeline in which DIAMOND alignment is performed against the AnnoTree protein database and we refer to this as an “AnnoTree run”. To enable the pipeline to be run in this way, we provide (1) a FastA file containing all ≈ 106 million AnnoTree protein sequences, and (2) an SQLite database (https://www.sqlite.org) that contains mappings of AnnoTree protein accessions to taxonomic and functional classes in all supported classifications. In addition, we provide a set of Python scripts that can be used to update both the FastA file and the mapping database.

To allow a fair comparison between the two runs, when aligning against the NCBI-nr database, throughout this study, we only aligned against the prokaryotic entries of the NCBI-nr database.

### Performance on a mock community

For an initial comparison of the two ways of running the DIAMOND+MEGAN pipeline, we ran both on a set of 53, 654 PacBio shotgun reads from the MBARC-26 mock community of 23 bacterial and 3 archaeal strains (30). In Figure 1 we compare the two taxonomic profiles obtained from the NCBI-nr and AnnoTree runs with the published profile. Both computed profiles follow the published one quite closely, with some low-abundance false-negative assignments, namely to *Salmonella bongori* and *Escherichia coli* for the former and latter run, respectively, and to *Nocardiopsis dassonvillei* for both. The NCBI-nr run gives rise to five false-positive bacterial assignments, namely to *Paenibacillaceae bacterium*, *Pseudomonas aeruginosa*, *Sediminispirochaeta bajacaliforniensis*, *Meiothermus hypogaeus* and *Thermobacillus sp.*. The latter has a large count that bleeds over from the high-abundance true positive *Thermobacillus composti*. The AnnoTree run gives rise to two false positive bacteria, *Sediminispirochaeta bajacaliforniensis* and *Meiothermus sp. Pnk-1*. There are no false positives or negatives in the archaea.

**FIG 1.**
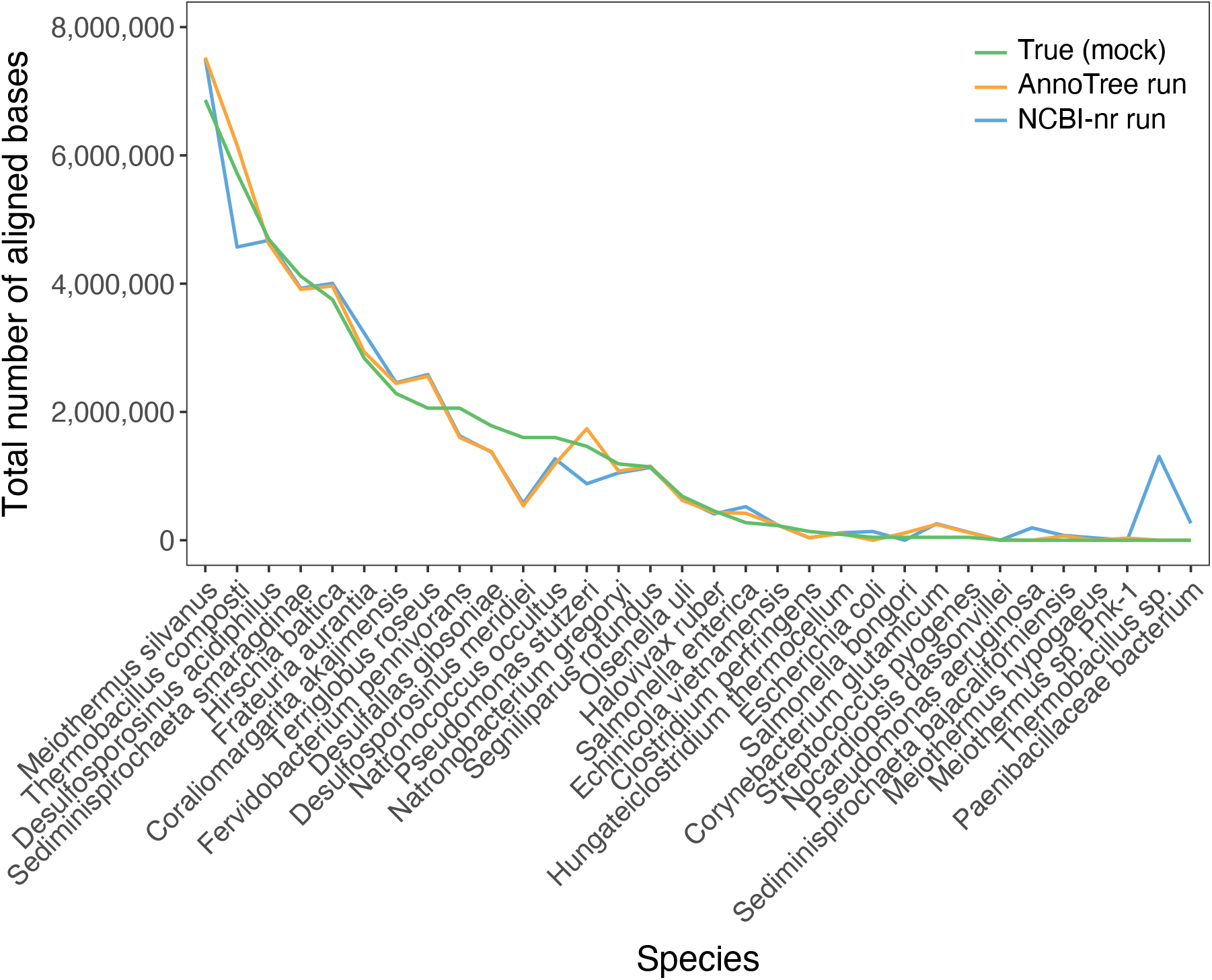
For prokaryotic species detected during analysis, we report the approximate relative abundance of organisms in the mock community (green), the number of bases aligned and assigned by DIAMOND+MEGAN in an NCBI-nr run (blue), and in an AnnoTree run (orange), respectively.

These results indicate that the use of the AnnoTree protein database, in place of the NCBI-nr protein database, is worth pursing when the goal is to determine the prokaryotic content of a sample.

### Performance on ten different datasets

For a more detailed comparison of NCBI-nr and AnnoTree runs of the DIAMOND+MEGAN pipeline, we applied the two variants to a set of ten different published samples that cover a range of different environments. Nine of the ten datasets consist of short reads and one consists of long reads. There are ≈ 580 million reads in total. The number of reads per dataset is listed in Table 1. For each dataset, we also report the number and percentage of reads that obtained a DIAMOND alignment against AnnoTree and NCBI-nr, respectively (Spearman’s correlation *ρ* = 1, *p* = 2.2*e* − 16). In most cases, the ratio of the former number to the latter is ≈ 1, except in the case of the Soil sample, where ≈ 50% more reads have an alignment against AnnoTree.

**Table 1.**
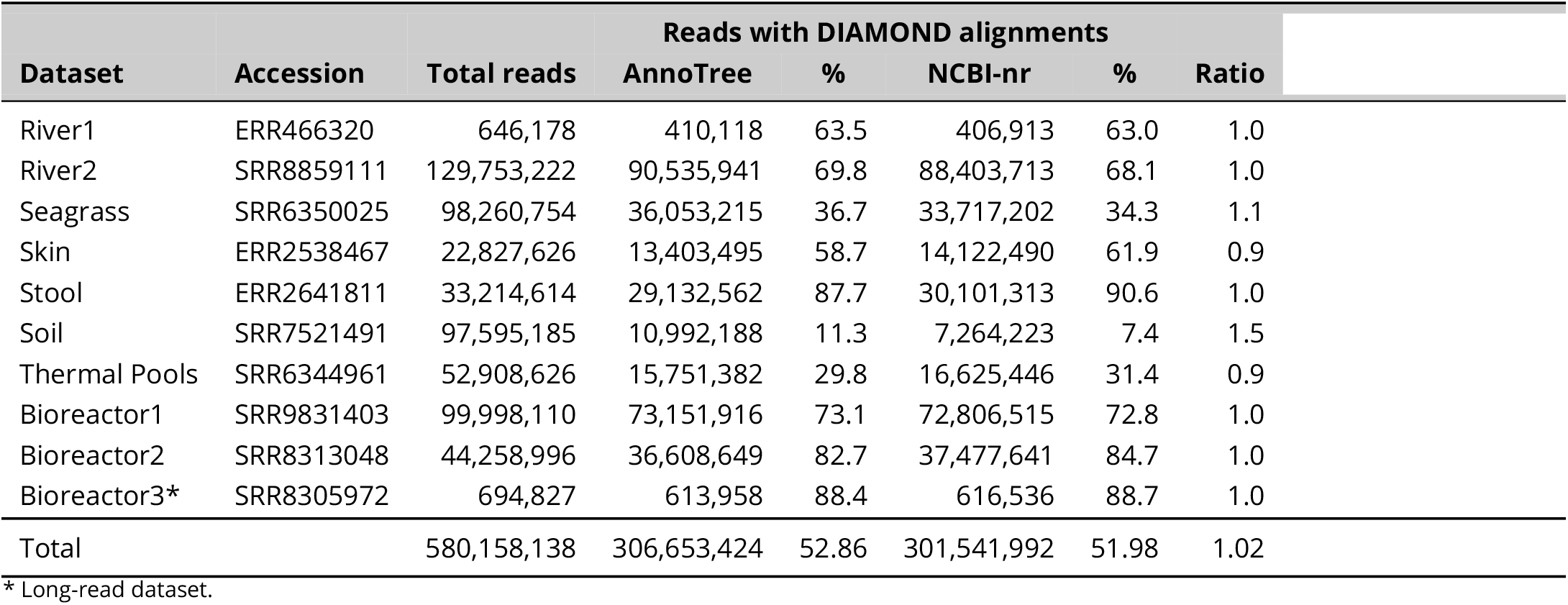
For each of ten datasets, we report the accession and total number of reads. In addition, for both the AnnoTree and NCBI-nr protein databases, we report the number and percentage of reads that obtained an alignment using DIAMOND, respectively. We also report the ratio between the two numbers.

| Dataset | Accession | Total reads | Reads with DIAMOND alignments |  |  |  | Ratio |
| --- | --- | --- | --- | --- | --- | --- | --- |
|  |  |  | AnnoTree | % | NCBI-nr | % |  |
| River1 | ERR466320 | 646,178 | 410,118 | 63.5 | 406,913 | 63.0 | 1.0 |
| River2 | SRR8859111 | 129,753,222 | 90,535,941 | 69.8 | 88,403,713 | 68.1 | 1.0 |
| Seagrass | SRR6350025 | 98,260,754 | 36,053,215 | 36.7 | 33,717,202 | 34.3 | 1.1 |
| Skin | ERR2538467 | 22,827,626 | 13,403,495 | 58.7 | 14,122,490 | 61.9 | 0.9 |
| Stool | ERR2641811 | 33,214,614 | 29,132,562 | 87.7 | 30,101,313 | 90.6 | 1.0 |
| Soil | SRR7521491 | 97,595,185 | 10,992,188 | 11.3 | 7,264,223 | 7.4 | 1.5 |
| Thermal Pools | SRR6344961 | 52,908,626 | 15,751,382 | 29.8 | 16,625,446 | 31.4 | 0.9 |
| Bioreactor1 | SRR9831403 | 99,998,110 | 73,151,916 | 73.1 | 72,806,515 | 72.8 | 1.0 |
| Bioreactor2 | SRR8313048 | 44,258,996 | 36,608,649 | 82.7 | 37,477,641 | 84.7 | 1.0 |
| Bioreactor3* | SRR8305972 | 694,827 | 613,958 | 88.4 | 616,536 | 88.7 | 1.0 |
| Total |  | 580,158,138 | 306,653,424 | 52.86 | 301,541,992 | 51.98 | 1.02 |

\* Long-read dataset.

Out of the total of ≈ 580 million reads, DIAMOND aligns ≈ 307 million (≈ 52.9%) reads against the AnnoTree database and ≈ 302 million (≈ 52%) reads against the NCBI-nr database, that is, nearly one percent more reads against the former database, although the latter database contains more than twice as many entries.

### Taxonomic binning

The DIAMOND+MEGAN pipeline assigns aligned reads both to the NCBI taxonomy (18) and, now, also to the GTDB taxonomy (19).

In the case of the NCBI taxonomy, performing AnnoTree runs on all datasets assigns 99.5% of all aligned reads to a taxonomic node, whereas the assignment rate when performing NCBI-nr runs is only 98.7%, see Table 2. In total, using AnnoTree rather than NCBI-nr leads to the taxonomic assignment of ≈ 7.6 million additional reads (≈ 1.3% of all reads). However, in more detail, the number of assigned reads is a few percent higher for the NCBI-nr runs on the two human-associated samples, Skin and Stool, and a few percent lower for the Seagrass and Soil samples.

**Table 2.**
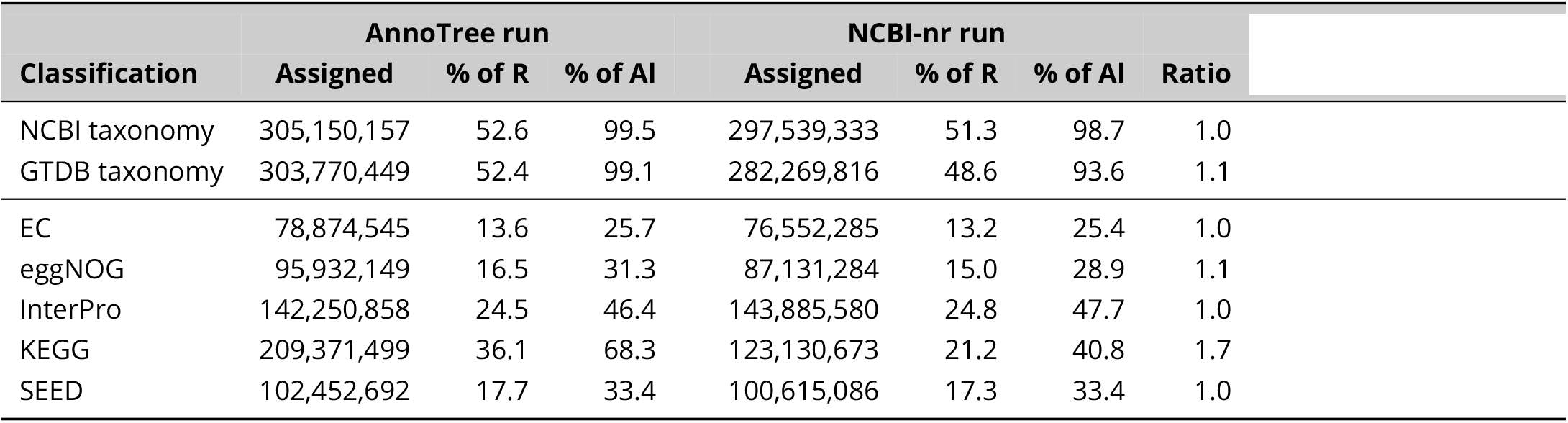
For each of the classifications provided by MEGAN, and summarized over all AnnoTree runs and NCBI-nr runs of the DIAMOND+MEGAN pipeline on the ten datasets listed in Table 1, we report the number of assigned reads (“Assigned”), also as a percentage of all reads (“% of R”), and as a percentage of all aligned reads (“% of Al”). In the last column, we report the ratio of the reads assigned using AnnoTree and using NCBI-nr, respectively.

| Classification | AnnoTree run |  |  | NCBI-nr run |  |  | Ratio |
| --- | --- | --- | --- | --- | --- | --- | --- |
|  | Assigned | % of R | % of AI | Assigned | % of R | % of AI |  |
| NCBI taxonomy | 305,150,157 | 52.6 | 99.5 | 297,539,333 | 51.3 | 98.7 | 1.0 |
| GTDB taxonomy | 303,770,449 | 52.4 | 99.1 | 282,269,816 | 48.6 | 93.6 | 1.1 |
| EC | 78,874,545 | 13.6 | 25.7 | 76,552,285 | 13.2 | 25.4 | 1.0 |
| eggNOG | 95,932,149 | 16.5 | 31.3 | 87,131,284 | 15.0 | 28.9 | 1.1 |
| InterPro | 142,250,858 | 24.5 | 46.4 | 143,885,580 | 24.8 | 47.7 | 1.0 |
| KEGG | 209,371,499 | 36.1 | 68.3 | 123,130,673 | 21.2 | 40.8 | 1.7 |
| SEED | 102,452,692 | 17.7 | 33.4 | 100,615,086 | 17.3 | 33.4 | 1.0 |

For each of the ten datasets, in Figure 2 we present a more detailed comparison of the assignment of reads to the NCBI taxonomy for the AnnoTree and NCBI-nr runs. The results for both runs agree for 30-60% of all reads. For most datasets, the percentage of assigned reads that are assigned by only one of the two runs is similar, the biggest exception being the Soil sample, where approximately 40% of all assigned reads are assigned by AnnoTree only. For most datasets, the percentage of reads that are assigned incompatibly to different lineages is below or around 10%, with the exception of a bioreactor sample, where this value is around 30%.

**FIG 2.**
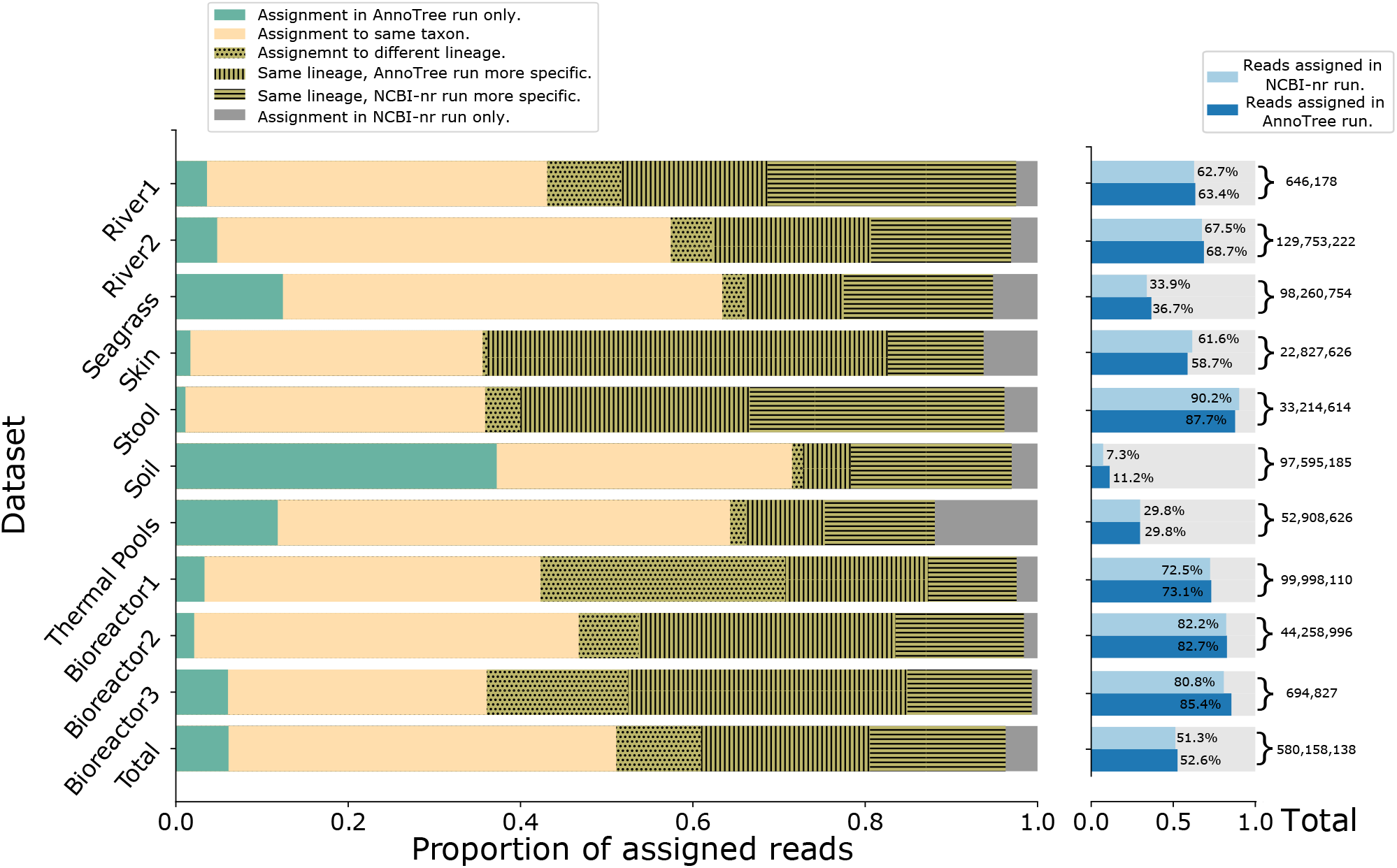
Details of the assignment of reads to the NCBI taxonomy. For each of the ten datasets, for the total set of reads assigned by either an AnnoTree run or NCBI run of the DIAMOND+MEGAN pipeline, we show the proportion of reads only assigned by the AnnoTree run (green), or assigned by both runs to the same taxon (yellow), or only assigned by the NCBI run (gray). For reads with differing assignments, we show the proportion assigned to incompatible lineages (dotted) or two compatible lineages with either the AnnoTree assignment being more specific (vertical stripes) or the NCBI-nr assignment being more specific (horizontal stripes). On the right, we indicate the total number of reads, and the number of reads assigned by either the AnnoTree or NCBI-nr run.

In the case of the GTDB taxonomy, performing AnnoTree runs on all datasets assigns ≈ 99% of all aligned reads to a taxonomic node, whereas the assignment rate when performing NCBI-nr runs is only 93.6%, see Table 2. In total, using AnnoTree rather than NCBI-nr leads to an assignment of ≈ 21.5 million additional reads (≈ 3.7% of all reads). On the stool sample, the assignment rate for the NCBI-nr run is one percent higher than for the AnnoTree run, but in all other cases, the assignment rate for the AnnoTree runs is a couple of percent higher than for the NCBI-nr runs.

In Figure 3, We present more details on the assignment of reads to the GTDB taxonomy. The results are similar to those for the assignment to the NCBI taxonomy, but with a somewhat decreased level of conflicting assignments for most datasets.

**FIG 3.**
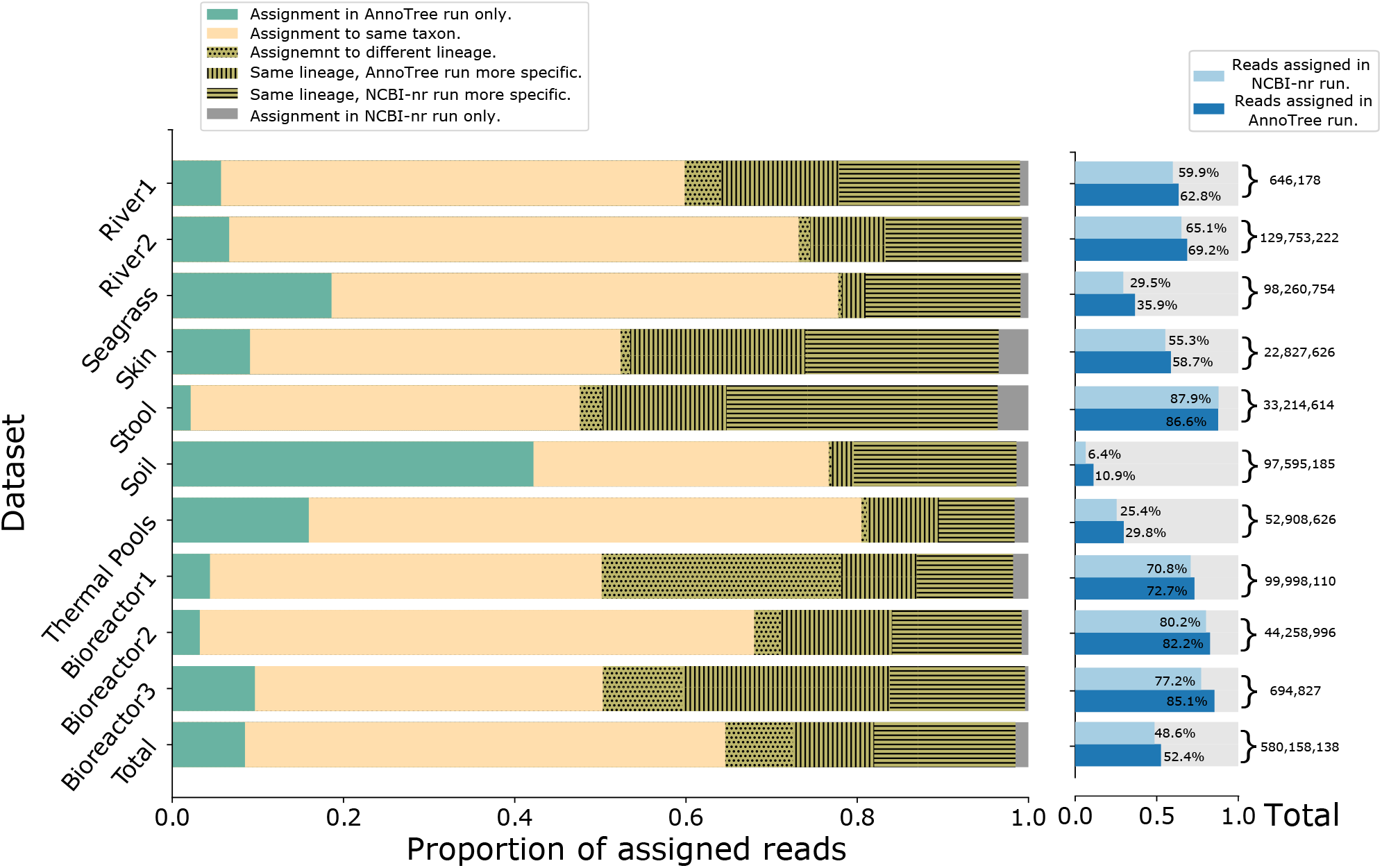
Details of the assignment of reads to the GTDB taxonomy, using the same colors etc as in the previous Figure.

### Functional binning

The DIAMOND+MEGAN pipeline assigns aligned reads to a number of different functional classifications, currently EC numbers (20), eggNOG (21), InterPro families (22), KEGG (23) and SEED (24). For most functional classifications, the assignment rates for the AnnoTree runs are slightly higher than for the NCBI-nr runs.

In the case of KEGG, performing AnnoTree runs on all datasets assigns ≈ 68.3% of all aligned reads to a KEGG node, whereas the assignment rate when performing NCBI-nr runs is only ≈ 40.8%, see Table 2. AnnoTree runs make ≈ 70% more assignments of reads to KEGG nodes than the NCBI-nr runs do.

The number of read assignments to KEGG is particularly low for the Soil sample, with AnnoTree-based assignment of only ≈ 8% and NCBI-based assignment of only ≈ 1.6% of all reads (see Figure 4).

**FIG 4.**
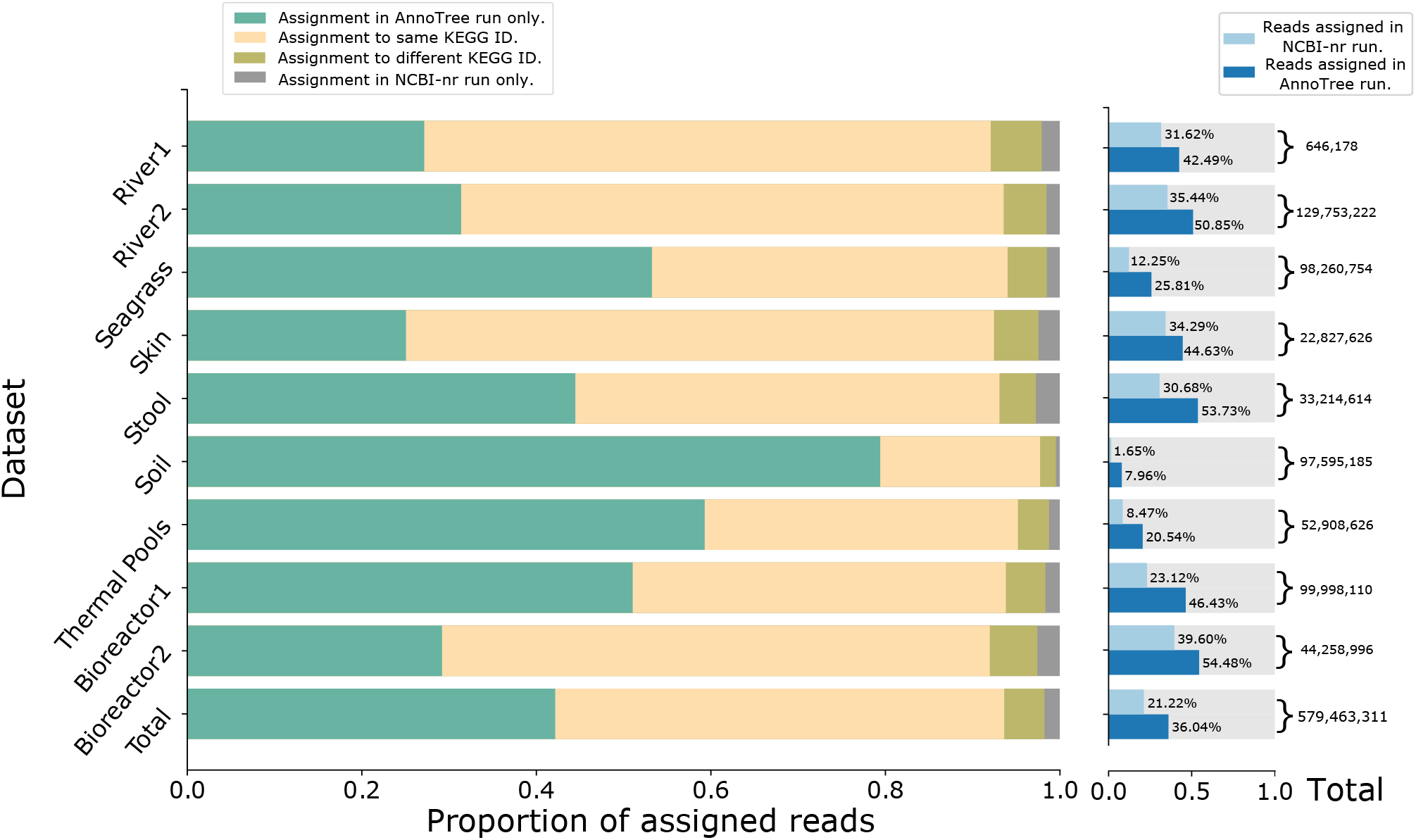
Details of the assignment of reads to KEGG. For each of the ten datasets, for the total set of reads assigned by either an AnnoTree run or NCBI run of the DIAMOND+MEGAN pipeline, we show the proportion of reads only assigned by the AnnoTree run (green), or assigned by both runs to the same class (yellow), or different classes (olive) or only assigned by the NCBI run (gray).

### Running time

As the AnnoTree protein database is less than one quarter the size of the full NCBI-nr protein database, using the former during the alignment step of the DIAMOND+MEGAN pipeline will speed up the analysis. This will be offset, very slightly, by the fact that the number of aligned reads will be slightly higher. In the analysis performed here, we only aligned against the *prokaryotic* content of NCBI-nr, which contains approximately twice as many sequences as the AnnoTree protein database.

In Table 3, summarizing over all ten datasets, we report the CPU time used by DIAMOND, Meganizer, and both combined, during both a NCBI-nr run and an AnnoTree run of the DIAMOND+MEGAN pipeline. On all datasets, DIAMOND alignment against the prokaryotic proteins in NCBI-nr takes about twice as along as alignment against the AnnoTree database, while meganization usually takes slightly longer in AnnoTree runs. In total, an AnnoTree run of the DIAMOND+MEGAN pipeline is twice as fast as an NCBI-nr run.

**Table 3.**
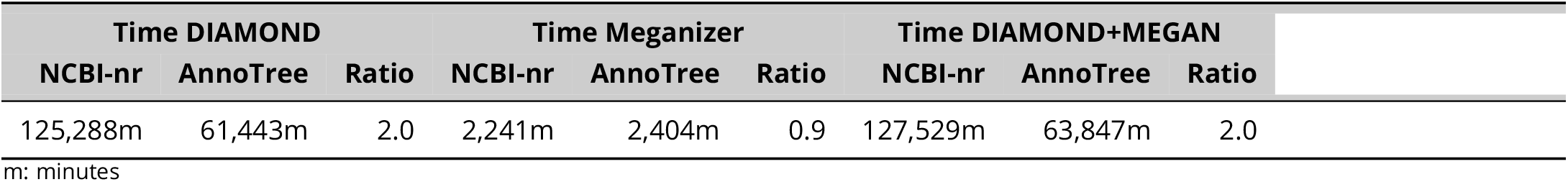
Summarizing over all ten datasets, we show the CPU time used for running DIAMOND, Meganizer, and both combined, during either an NCBI-nr run (restricted to prokaryotic sequences) or an AnnoTree run, of the DIAMOND+MEGAN pipeline, respectively.

Performing an NCBI run using the *full* NCBI-nr database, not just the prokaryotic part, on each of the ten datasets, takes ≈ 3 times as long as an AnnoTree run, with a ≈ 4% increase of assignments to the NCBI taxonomy, on average.

## DISCUSSION

When first published in 2007 (10), MEGAN was run together with BLASTX (9) on datasets containing hundreds of thousands of short reads against the NCBI-nr protein database, which then contained around 3 million sequences. The DIAMOND alignment program (14) was later designed to allow the alignment of much larger datasets against a much larger NCBI-nr database. As the NCBI-nr database continues to grow, alignment against the full database presents an increasingly severe computational bottleneck.

As an alternative, projects focusing on the prokaryotic content of microbiome samples can make use of the GTDB taxonomy and the AnnoTree protein database, which are both explicitly designed for this. As the AnnoTree protein database is only 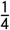 the size of the NCBI-nr database, DIAMOND alignment of reads against the AnnoTree protein database takes at most only half as much time, while providing a superior assignment rate.

In this study, we illustrated the performance of AnnoTree runs using ten example datasets, from different environments. The results on these examples confirm that using AnnoTree is a useful alternative to using NCBI-nr.

The Soil sample stands out. Its DIAMOND-alignment rates against AnnoTree and NCBI-nr are very low at only 11.3% and 7.4%, respectively, in contrast to the alignment rates for the Stool sample, say, which are very high at 87.7% and 90.6%, respectively. This illustrates that the diversity of the soil environment is only poorly represented in current databases (31, 32), while human stool samples and human-associated microbes have been studied in detail (33). The fact that the AnnoTree run has a higher assignment rate than the NCBI-nr run on the Soil sample may be due to the fact that the AnnoTree database recruits more sequences from “metagenomic assembled genomes” than NCBI-nr does.

To extend the AnnoTree-based approach to the detection and analysis of viral sequences, one could extend the AnnoTree protein database using either the virus subdivision of NCBI-nr or another dedicated resource (34, 35).

## MATERIALS AND METHODS

### Datasets

We downloaded 53, 654 PacBio shotgun reads (SRA accession SRR3656744) from the MBARC-26 (Mock Bacteria ARchaea Community) dataset (30), length 11 - 16, 403, mean 1, 643.5. The true community profile reported in Figure 1 was estimated from Figure 2a of (36).

The ten example datasets, listed in Table 4, were downloaded in FASTA format from the NCBI Sequence Read Archive (SRA) using the NCBI SRA toolkit’s fastq-dump program, as follows:

**Table 4.** For each of the ten datasets, we list their SRA Run Id, sequencing platform, read layout and total number reads.

| Dataset | SRA Run Id | Platform | Layout | Total Reads |
| --- | --- | --- | --- | --- |
| River1 | ERR466320 | LS454 | single | 646,178 |
| River2 | SRR8859111 | Illumina | paired | 129,753,222 |
| Seagrass | SRR6350025 | Illumina | paired | 98,260,754 |
| Skin | ERR2538467 | Illumina | single | 22,827,626 |
| Stool | ERR2641811 | Illumina | paired | 33,214,614 |
| Soil | SRR7521491 | Illumina | paired | 97,595,185 |
| Thermal Pools | SRR6344961 | Illumina | paired | 52,908,626 |
| Bioreactor1 | SRR9831403 | Illumina | paired | 99,998,110 |
| Bioreactor2 | SRR8313048 | Illumina | paired | 44,258,996 |
| Bioreactor3 | SRR8305972 | ONT | single | 694,827 |

fastq-dump –split-spot –fasta 80 -I accession

Datasets with paired-end reads were concatenated into a single file. No additional preprocessing was performed.

In more detail, we used two datasets from rivers, River1 https://www.ncbi.nlm.nih.gov/sra/?term=ERR466320 and River2 (37), one from the seagrass rhizosphere (38), one from the skin (39), one from the stool (40), one from the soil (41), one from thermal pools (42), and three from bioreactors (Bioreactor1 (43), Bioreactor2 (44) and Bioreactor3 (44)). Nine of the ten datasets consist of short reads, whereas the last dataset consists of ONT MinION long reads.

### Protein reference databases

The NCBI-nr protein database was downloaded in January 2021 from the NCBI FTP site using the link ftp://ftp.ncbi.nlm.nih.gov/blast/db/FASTA/nr.gz. We also downloaded the two files prot.accession2taxid.gz ftp://ftp.ncbi.nlm.nih.gov/pub/taxonomy/accession2taxid/prot.accession2taxid.gz and nodes.dmp ftp://ftp.ncbi.nlm.nih.gov/pub/taxonomy/taxdmp.zip.

A DIAMOND index was then generated using the following command (requiring 150 CPU minutes):

diamond makedb -i nr.gz -d nr --taxonmap prot.accession2taxid.gz --taxonnodes nodes.dmp

For both the AnnoTree Bacteria database and the AnnoTree Archaea database, we downloaded MySQL dump files (version of Aug 25, 2020) from the AnnoTree Bitbucket repository https://bitbucket.org/doxeylabcrew/annotree-database/src/master/. These files were then imported into a MySQL server (version 5.7.35). Both databases each have 21 tables, which hold information on the AnnoTree hierarchy, the GTDB taxonomy, the NCBI taxonomy, protein sequences as well as additional mappings to Pfam, TIGRFAMs and KEGG.

For each sequence in the ‘protein_sequences’ table, we constructed a unique two-part accession string by concatenating its ‘gene_id’ and ‘gtdb_id’ values. For example, the protein sequence with gene ID AE009439_1_1 and GTDB genome ID GB_GCA_000007185_1 was given the two-part accession AE009439_1_1 GB_GCA_000007185_1.

These accessions and the corresponding protein sequences (from both databases) were written to a FastA file annotree.fasta, which we make available here: https://software-ab.informatik.uni-tuebingen.de/download/megan-annotree.

A DIAMOND index was then generated using the following command (requiring 33 CPU minutes):

diamond makedb -i annotree.fasta.gz -d annotree

### MEGAN mapping databases

We use the term “meganization” to refer to the process of analysing the alignments of a set of sequences so as to perform taxonomic binning (for example, using the naïve LCA algorithm for short reads or the interval-union LCA for long reads) and functional binning (usually using a best-hit approach). Meganization of DIAMOND alignments is performed either interactively using MEGAN or in a commandline fashion using the daa-meganizer program, which is bundled with MEGAN.

To perform meganization, MEGAN requires a so-called mapping database. This is an SQLite database file that contains a mapping of protein sequence accessions to all used taxonomical and functional classifications, namely the NCBI taxonomy, the GTDB taxonomy, EC, EGGNOG, INTERPRO families, KEGG (MEGAN Ultimate Edition) and SEED. For NCBI-nr runs, we used the mapping database megan-map-Jul2020-2-ue.db, which we downloaded from https://software-ab.informatik.uni-tuebingen.de/download/megan6.

For AnnoTree runs, we created a new mapping database called megan-mapping-annotree-June-2021.db in SQLite format. This file is available here: https://software-ab.informatik.uni-tuebingen.de/download/megan-annotree

We used the above described two-part accessions as the primary key for the ‘mapping’ table. We determined the other entries of the ‘mapping’ table as follows. The value for the GTDB and NCBI taxonomies were obtained from the ‘node_tax’ tables of the two MySQL databases described above, using the gtdb_id part of the two-part accession.

The value for the KEGG classification was obtained from the ‘kegg_top_hits’ tables of the two MySQL databases, using the gene_id part of the two-part accession. In the case that there is more than one possible KEGG assignment for a given protein, we randomly selected one. This was necessary because MEGAN allows only at most one assignment per reference sequence. Both GTDB and KEGG IDs were additionally formatted to match the format required by MEGAN.

We calculated entries for the other classifications supported by MEGAN (EC,EGGNOG, INTERPRO and SEED) by performing a join on the MD5 hash values of the protein sequences in the NCBI-nr and AnnoTree protein databases, in other words, by copying the classifications of an NCBI-nr accession over to an AnnoTree two-part accession, whenever the two accessions correspond to exactly the same protein sequence. We list the number of accessions that have assignments in the different classifications in Table 5.

**Table 5.** For two different taxonomical classifications (NCBI and GTDB), and for five different functional classifications (EC, EGG, InterPro, KEGG and SEED) supported by MEGAN, we report the number of prokaryotic accessions in NCBI-nr or AnnoTree, that have a mapping to some class in the classification, and the corresponding ratio.

| Classification | NCBI-nr | AnnoTree | Ratio |
| --- | --- | --- | --- |
| NCBI taxonomy | 182,329,414 | 106,052,079 | 1.72 |
| GTDB taxonomy | 126,956,422 | 106,052,079 | 1.2 |
| EC | 4,501,593 | 2,962,187 | 1.51 |
| eggNOG | 4,274,800 | 3,506,041 | 1.21 |
| InterPro | 19,748,423 | 11,069,757 | 1.78 |
| KEGG | 8,218,708* | 56,577,432 | 0.15 |
| SEED | 31,117,272 | 16,183,436 | 1.92 |

\*MEGAN Ultimate Edition.

For most functional classifications, the number of AnnoTree proteins with assignments is smaller than for NCBI-nr proteins, which is due to the fact that the AnnoTree assignments are copied from NCBI-nr assignments. In the case of KEGG, the assignments are obtained from the AnnoTree database, which appears to maintain a much richer mapping than previously provided by MEGAN.

### Dataset processing

We ran DIAMOND (v2.0.6.144) on each short-read dataset as follows:

diamond blastx -d 〈index〉 -q 〈input〉 -o 〈output〉 -f 100 -b24 -c1

Here, 〈index〉 is the index file, either annotree or nr, computed as described above. The input file, in (compressed) FastA or FastQ format, is specified by 〈input〉 and the output file is specified by 〈output〉. We used -f 100 to specify output in DAA format. The two remaining options -b24 -c1 were used in an attempt to tune performance.

In addition, for purposes of comparison against the AnnoTree database, when running DIAMOND on NCBI-nr, we used the option --taxonlist 2,2157 to restrict alignment to bacteria (taxon ID 2) and archaea (taxon ID 2157).

When processing long reads, we also specified the --long-reads option.

The resulting DAA files were meganized using the daa-meganizer program MEGAN (version 6.21.5, Ultimate Edition, built 5 May 2021), as follows:

tools/daa-meganizer -i 〈input〉 -mdb 〈mapping〉

Here, the input file 〈input〉 is a DAA file produced by DIAMOND and the mapping file 〈mapping〉 was either megan-map-Jul2020-2-ue.db or megan-mapping-annotree-June-2021.db, depending on whether the DIAMOND run was against the NCBI-nr or AnnTree protein database, respectively. When processing long reads, we also specified the -lg option.

### Data comparison

We used the MEGAN tool daa2info to extract the mapping of reads to taxonomic and functional classes obtained in both the NCBI-nr and AnnoTree runs of the DIAMOND+MEGAN pipeline.

The following command was used to extract the mapping of reads to classes for all classifications:

tools/daa2info -i 〈input〉 -o 〈output〉 -l -m -r2c Taxonomy GTDB KEGG EC EGGNOG INTERPRO2GO SEED

Here, the input file 〈input〉 is a meganized DAA file and the output file 〈output〉 is a text file. The output was used to determine the assignment rates for different classifications.

Similarly, the following command was used to extract the mapping of reads to taxonomic paths in the NCBI taxonomy:

tools/daa2info -i 〈input〉 -o 〈output〉 -l -m -r2c Taxonomy -p true -r true

The output was used to generate Figure 2.

The following command was used to extract the mapping of reads to taxonomic paths in the GTDB taxonomy:

tools/daa2info -i 〈input〉 -o 〈output〉 -l -m -r2c GTDB -p true -r true.

The output was used to generate Figure 3.

### Computational resources

The DIAMOND+MEGAN pipeline was run on a linux server with 28 cores, 512GB of RAM and 2TB of disk. Reported run-times are “user time” as calculated using the linux “time” command. All other calculations where undertaken on a MacBook Pro latop with 12 cores and 16GB RAM.

### Statistical analysis

Spearman’s correlations were computed using ggplot2 (45).

### Availability of data and materials

All datasets analyzed here are publicly available from NCBI SRA, using the accessions listed in Table 4.

The AnnoTree protein FastA file and mapping database can both be downloaded from https://software-ab.informatik.uni-tuebingen.de/download/megan-annotree.

## ACKNOWLEDGMENTS

This work was supported by the BMBF-funded de.NBI Cloud within the German Network for Bioinformatics Infrastructure (de.NBI) (031A532B, 031A533A, 031A533B, 031A534A, 031A535A, 031A537A, 031A537B, 031A537C, 031A537D, 031A538A).

The authors would also like to acknowledge support by the High Performance and Cloud Computing Group at the Zentrum für Datenverarbeitung of the University of Tübingen, the state of Baden-Württemberg through bwHPC and the German Research Foundation (DFG) through grant no INST 37/935-1 FUGG.

The authors acknowledge infrastructural support by the cluster of Excellence EXC2124 Controlling Microbes to Fight Infection (CMFI), project ID 390838134.

CB was supported by the German Research Foundation (DFG) through grant no HU 566/12-1.

Furthermore, we acknowledge support by the Open Access Publishing Fund of University of Tübingen.

## AUTHORS’ CONTRIBUTIONS

DHH and CB conceptualized the project. HF and CB performed the computations. AG and HF analyzed the results. AG and DHH wrote the manuscript. All authors edited the manuscript.

